# MGM as a large-scale pretrained foundation model for microbiome analyses in diverse contexts

**DOI:** 10.1101/2024.12.30.630825

**Authors:** Haohong Zhang, Yuli Zhang, Zixin Kang, Lulu Song, Ronghua Yang, Kang Ning

## Abstract

Microbial communities significantly impact medicine, biotechnology, and agriculture. Advanced sequencing technologies have generated extensive microbiome data, enabling the discovery of substantial evolutionary and ecological patterns. However, traditional supervised learning methods struggle to capture universal patterns in microbial community data, largely due to the large data heterogeneity and profound batch effects among samples, rendering it difficult to classify samples as well as detect biomarkers from millions of samples, not to say the intricate but important dynamic patterns from a variety of contextualized sceneries. In this study, we introduce the Microbial General Model (MGM), the first microbiome community foundation model pre-trained on a dataset of 263,302 microbiome samples using language modeling techniques. MGM demonstrated significant improvements in microbial community classification compared to traditional machine learning methods. Additionally, MGM has enabled contextualized classification, effectively overcomes cross-regional limitations, showing enhanced performance on intercontinental datasets through transfer learning. Furthermore, fine-tuning MGM on a longitudinal infant dataset revealed distinct keystone genera during development, with *Bacteroides* and *Bifidobacterium* exhibiting higher attention weights in vaginal deliveries, and *Haemophilus* in cesarean deliveries. Finally, through in silico modeling, the model also uncovered novel microbial dynamic patterns in a Crohn’s disease cohort following antibiotic treatment. In conclusion, by leveraging self-attention and autoregressive pre-training, MGM serves as a versatile model for various downstream microbiome tasks and holds significant potential for achieving contextualized aims.

**Key points:**

- The Microbial General Model (MGM) is a foundation model with millions of parameters pre-trained on sub-million microbial community data.
- MGM outperforms traditional methods in various microbiome classification and prediction tasks, such as microbial community classification.
- MGM effectively captures the spatial and temporal dynamics of microbial communities.
- MGM could detect the effects of perturbation on microbial community through in silico experiments.

## Introduction

Microbial communities, ubiquitous across diverse environments play crucial roles in shaping ecological niches have significant implications for health [1, 2], synthetic biology [3, 4], and environmental science [5, 6]. The advent of sequencing technologies has enabled researchers to amass vast microbiome datasets, significantly expanding our understanding of these complex systems [7]. Currently, hundreds of thousands of microbiome samples and their sequencing data have been accumulated and deposited in public databases [8]. For example, as of 2023, EBI MGnify, a leading platform for microbiome data analysis and archiving, has cataloged 343,695 distinct samples from 4,601 studies across various biomes, including environmental, engineered, and host-associated microbiomes [8]. While these extensive collections of microbial community samples represent a valuable resource, they also present challenges in integrating large-scale microbiome data and extracting complex, multifaceted patterns within microbial communities, which is essential for advancing our understanding of their subtle evolutionary and ecological dynamics [9]. However, data heterogeneity, including insufficient data standards and a lack of interoperability across datasets, as well as profound batch effects among studies, limits the ability of traditional meta-analytical methods in capturing shared insights across studies [10–12].

A promising approach for overcoming these limitations is the use of foundation models. These models are pre-trained on large-scale datasets, enabling them to generate a broad range of outputs. Recently, several studies have focused on developing foundation model to improve the understanding of microbial sequence data [13–15]. These methods derive foundation models by training on vast, diverse datasets, capturing broad patterns that represent generalized biological relationships. For example, gLM, a genomic language model pre-trained on millions of metagenomic scaffolds, demonstrated impressive zero-shot performance in downstream tasks [15].

As the foundation model learned the shared knowledge embedded in large-scale dataset, it can be transferred to specific downstream tasks with the specific characteristics and nuances by transfer learning [16, 17]. This approach is often more effective than training models from scratch, as it leverages previously acquired knowledge, leading to improved performance and reduced training time [18–20]. By adjusting the learned representations to fit the context of the new dataset, transfer learning has shown potential for addressing the complexities of microbiome data integration and analysis [21–23].

However, these methods face inherent limitations due to their reliance on supervised learning strategies, which can introduce label bias during pre-training. Consequently, different downstream tasks often necessitate the adoption of distinct pre-trained models. Inappropriate pre-trained model selection can lead to either negligible performance gains or even performance degradation in the contextualized model [22]. Moreover, inaccurately annotated microbiome samples, such as those labeled “Mixed biome” in MGnify, can distort the training process, leading to misinterpretations and ultimately hindering model performance.

To address these limitations, self-supervised learning offers a compelling alternative. Unlike supervised methods, self-supervised learning does not rely on labeled data and instead allows models to uncover underlying patterns in large datasets [24]. This approach circumvents the need for high quality labeled datasets and has shown significant promise in natural language processing (NLP), where large language models (LLMs) have evolved from simple pattern recognition to tackling more sophisticated tasks such as reasoning and content generation [25]. A key driver of these capabilities is the self-attention mechanism [26], a core component of LLMs, which enables models to focus on relevant parts of high-dimensional input data, capturing contextual relationships within text. By fine-tuning these models on specific tasks, the knowledge gained during extensive pre-training on large corpora can be effectively transferred to new applications. This approach has led to significant performance improvements across a wide range of NLP tasks [27].

Recent advancements in the application of LLMs have shown promises in the analysis of biological tabular data [28–31]. Building on their success in processing textual data, these models have been adapted to handle the structured, high-dimensional nature of biological datasets. For example, scBERT employs a transformer-based architecture to analyze gene expression data, enhancing tasks such as cell type classification and gene expression imputation [28]. Geneformer leverages pre-trained LLMs to identify gene network regulatory elements within biological data [29]. Similarly, scGPT, inspired by generative pre-training, provides insights into cellular heterogeneity and simulating biological states under various conditions [30]. Lastly, scFoundation integrates foundation LLM techniques to create a versatile framework for diverse biological tasks, including cell type annotation, trajectory inference, and differential expression analysis [31]. These models underscore the potential of LLMs to revolutionize the analysis of biological tabular data, offering more accurate and comprehensive insights into complex biological systems. Building on these advancements in genomics analysis, the application of LLMs to the study of microbial communities offers a promising new frontier for understanding complex microbial ecosystems.

In this study, we introduce the Microbial General Model (MGM), a context-aware, attention-based foundation model specifically designed for microbiome analysis. MGM employs multi-layer transformer blocks and is pre-trained on nearly one million microbiome samples from diverse biomes to capture generalizable microbial composition patterns. Through transfer learning, MGM replaces its language modeling head with task-specific heads, enabling fine-tuning on limited data for various applications. In benchmark evaluations, MGM outperformed traditional machine learning approaches across multiple tasks. For example, in a cross-regional disease diagnosis task, MGM generalized effectively across diverse cohorts, overcoming intercontinental variations in microbial community structures. In a longitudinal infant cohort, MGM distinguished between delivery modes, identifying developmental distinctions and keystone species. In tumor microbiome analyses, MGM uncovered potential therapeutic targets through in silico perturbation experiments. Additionally, in a Crohn’s disease cohort, MGM revealed microbial dynamics influenced by antibiotic treatment, identifying consistent and novel microbial signals over time.

## Results

### Microcorpus-260K and MGM architecture

MGM is a foundation model pre-trained on large-scale microbiome community samples from various biomes. To facilitate this pre-training, we assembled Microcorpus-260K, a comprehensive dataset containing microbiome samples from the MGnify database up to June 2023. After filtering out low-quality or incomplete data, 263,302 samples were retained for pre-training (**Methods**). From these samples, we generated a vocabulary comprising 9,665 distinct genera. The genera were normalized and ranked based on their relative abundance in each sample, and then transformed into discrete input representations. Given the fixed input length required by the transformer model, we selected 512 as the input length. This choice was made to ensure that 99.99% of the samples were adequately covered without truncation (**Fig. 1a**). Following the preparation of the dataset, we developed MGM using a multi-layer transformer architecture (**Fig. 1c**), designed to effectively capture the patterns and structures present within the large-scale microbial community data.

**Figure 1.**
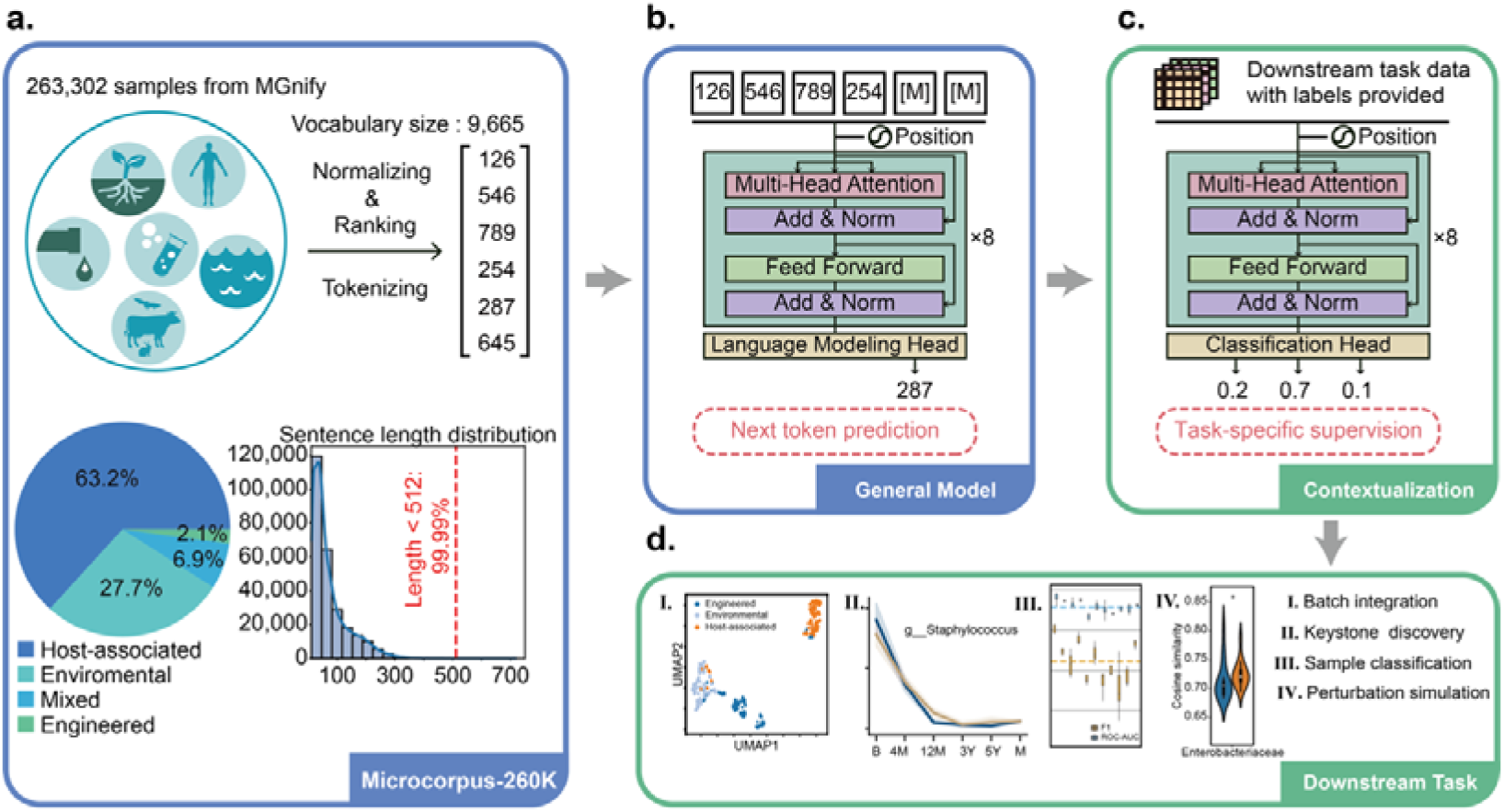
MGM architecture and transfer learning strategy. **a.** Construction process of Microcorpus-260K. **b.** MGM pre-trained using causal language modeling approach. **c.** Model details in multi-layer transformer blocks. **d.** Adaptation of MGM to different downstream tasks using transfer learning methods. **e.** Examples of downstream tasks: I. Batch integration based on contextualized sample embedding. II. Keystone species discovery based on contextualized attention weights. III. Accurate predictions based on transfer learning. IV. In silico perturbation analysis.

### Language modeling enables generalizable patterns capture

We pre-trained MGM using a causal language modeling approach on the Microcorpus-260K dataset. During this process, the model progressively learned to predict the next genera based on existing microbial composition within the sample (**Fig. 1b**). By leveraging the self-attention mechanism, MGM captures high-dimensional global representations of microbial community information (**Fig. 1c)**. For downstream tasks, MGM employs transfer learning strategy, where the language modeling head is replaced by task-specific heads (e.g., a sequence classification head) and the model is fine-tuned on limited data (**Fig. 1d**). This capacity makes it a versatile tool for a range of microbiome analyses, including batch integration, keystone species discovery and microbial community classification. Additionally, MGM can predict the effects of perturbation by modifying the rank of certain microorganisms (**Fig. 1e**).

To optimize model performance, we conducted a grid search on hyperparameters and selected a structure comprising 8 layers and 8 attention heads (**Fig. 2a, Supplementary Fig. 1a**). To evaluate the effectiveness of pre-training process, we randomly selected 1,000 samples from the validation set. For each sample, the pre-trained MGM model predicted microbial compositions (referred to as “sentences”) based on a proportion of genera (referred to as “tokens”). We then compared the cosine similarity between the embeddings of the predicted and original compositions. Remarkably, even when only 20% of the original genera were provided, the cosine similarity between the predicted and original embeddings exceeded 0.9 (**Fig. 2b**). These results highlighted MGM’s strong ability to capture and generalize microbiome patterns, positioning it as a powerful tool for various downstream applications.

**Figure 2.**
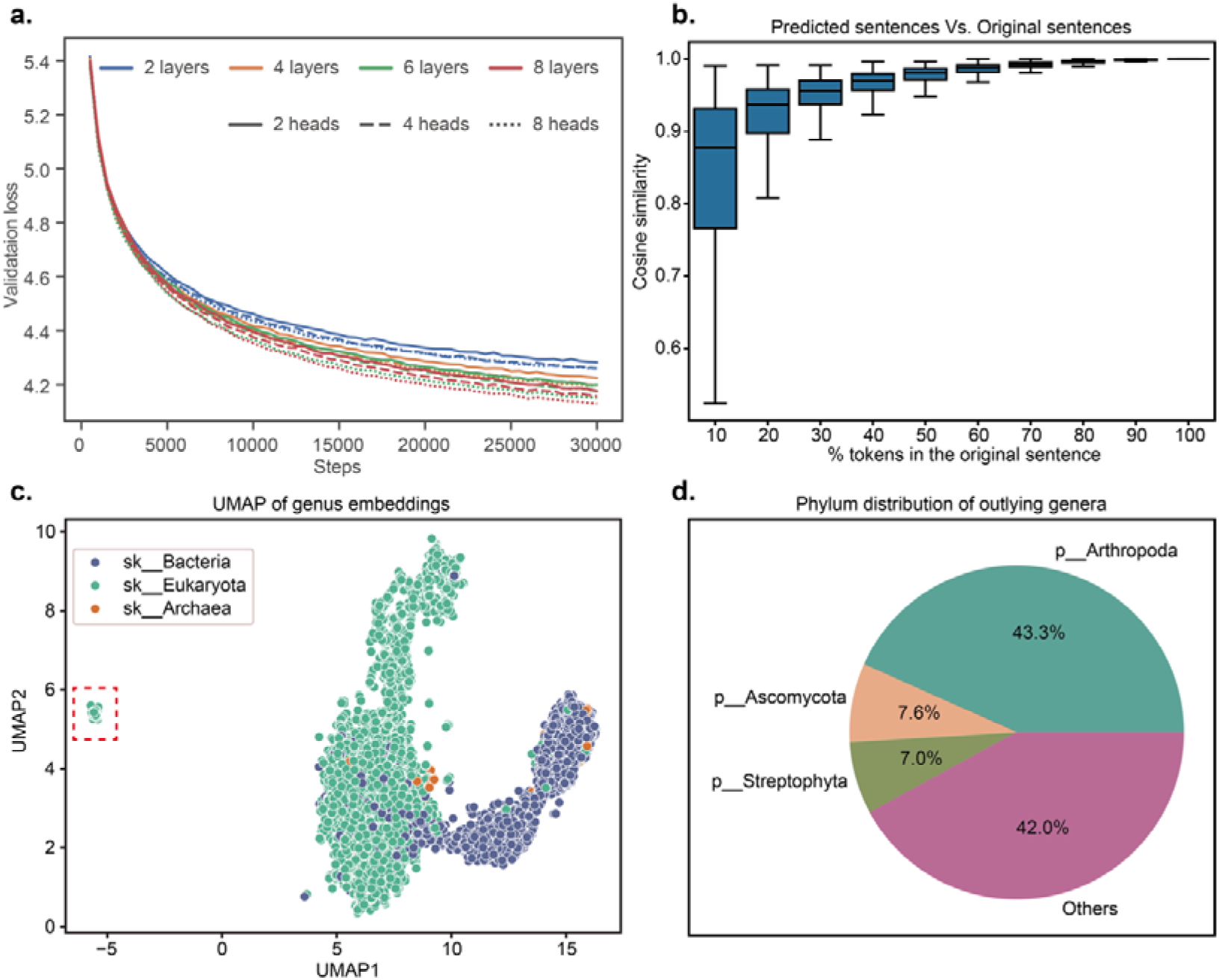
Hyperparameter search and pre-training evaluation. **a.** Grid search result of layers and heads of MGM. **b.** Similarity between generated sentences and the original sentences. **c.** UMAP visualization of the 9,665 genus embeddings extracted from MGM’s word embedding layer. **d.** Phylum-level distribution of the 157 genera identified as outliers in the UMAP plot. Only the top 3 phylum have the most genera are labeled.

We further explored whether the pre-training process captured taxonomic differences between genera, despite the absence of explicit phylogenetic information in our encoding strategy. We extracted embeddings for the 9,665 genera from the word embedding layer and found that genera from *Bacteria* and *Eukaryota* formed two diffuse clusters in the embedding space (**Fig. 2c)**. Additionally, we identified an outlier cluster comprising 157 genera, 43.3% of which belonged to *Arthropoda* (68 of 157, **Fig. 2d**). Given that *Arthropoda* is not typically considered a core component in microbiome studies, this finding suggested that MGM is capable of detecting taxonomic distinctions, even without explicit phylogenetic encoding.

### Microbial community classification and batch integration

We evaluated MGM by a comprehensive benchmark using a microbial community classification task on our Microcorpus-260K dataset. This task is a critical aspect of microbiome analysis, with applications including microbial source tracking [32] and noninvasive diagnostics [33]. We fine-tuned MGM for microbial community classification with cross-entropy on biome name and lineage from MGnify, followed by a 5-fold cross-validation on each biome layer. We compared these results with both traditional source tracking methods, such as FEAST [34], and other machine learning techniques, including K-Nearest Neighbor (KNN), Logistic Regression (LR), Random Forest (RF). To assess the value of self-supervised pre-training, we also evaluated an un-pre-trained MGM model. Our results demonstrated that the fine-tuned MGM significantly enhanced the ability to distinguish the source of samples (**Fig. 2a**, average ROC-AUC = 0.99), outperforming traditional methods (**Fig. 2a**, FEAST, average ROC-AUC = 0.68; KNN, average ROC-AUC = 0.94; LR, average ROC-AUC = 0.95; RF, average ROC-AUC = 0.97) and the un-pre-trained MGM (**Fig. 3a**, average ROC-AUC = 0.97). Notably, the EM-based FEAST method exhibited inefficiently, leading us to evaluate it only at the first layer. To further tested model generalization, we applied MGM on 43,528 new samples introduced to MGnify after March 2023 without additional fine-tuning. While the RF model performed slightly better in the shallower layers (layer 1 and 2), MGM excelled in the deeper, more complex layers (layer 3, 4, and 5) (**Fig. 3b**). Furthermore, both fine-tuned and un-pre-trained MGM embeddings yielded superior microbial community classification compared to community abundance profiles based on clustering performance (**Fig. 3c**). These findings underscored the value of the general patterns captured during self-supervised pre-training, which provided a foundational understanding of microbial community structures and enhanced downstream classification tasks.

**Figure 3.**
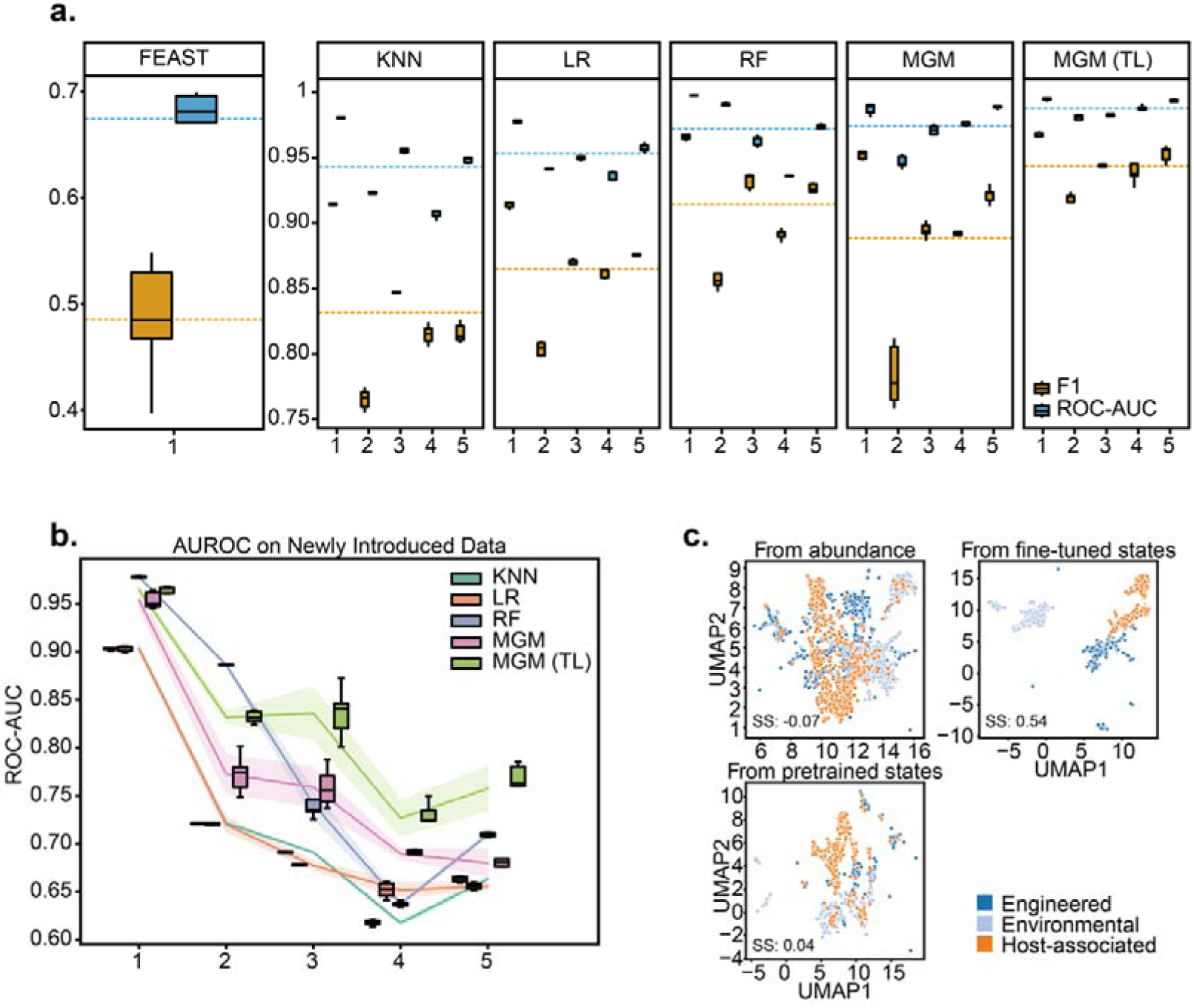
MGM Enhanced Source Tracking of Microbial Communities. **a.** Boxplot evaluation of the six methods using 5-fold cross-validation on each layer of biome lineage, with blue representing ROC-AUC, orange representing F1, and dashed lines indicating the average of all experiments. **b.** ROC-AUC performance of each method on newly introduced data in MGnify. We excluded FEAST for its poor computational inefficiency and performance at the shallow layer**. c.** UMAP dimensionality reduction of for 3,000 random samples from microbial relative abundance, pre-trained MGM embeddings and fine-tuned MGM embeddings colored by the biome of layer 1. KNN: K-nearest neighbors. LR: Logistic regression. RF: Random Forest. TL: transfer learning. SS: Silhouette score.

### Overcoming cross-regional limitation

Cross-regional diagnosis poses a significant challenge due to the variability in microbiome compositions across geographic regions [35]. Factors like diet, environment, and genetic background create distinct microbial profiles, complicating the development of diagnostic models that perform consistently across diverse populations. Traditional methods often struggle with overfitting to specific training data, which hinders their ability to generalize, especially when datasets are small or lack diversity. This variability highlights the need for a foundation model that can capture broad microbial patterns while being flexible enough to accommodate regional differences. MGM is pre-trained on large-scaled dataset, enabling it to recognize generalizable microbial features that are less susceptible to regional biases. When applied to cross-regional diagnosis, MGM can be fine-tuned with region-specific data, enhancing its ability to adapt to local microbial variations without losing the robustness of its general foundation.

We evaluated the robustness of MGM against regional limitations in clinical diagnosis. In such tasks, region-specific and disease-specific microbes often play a crucial role. Three models were evaluated for disease diagnosis performance across two intercontinental gut microbiome cohorts: an un-pre-trained model, a pre-trained model, and a Cross-country model fine-tuned with data from the target country.

For the inflammatory bowel disease (IBD) cohort [35] consisting of samples from Ireland and Canada, we observed that *Lachnoclostridium* ranks lower in the Crohn’s disease group compared to the healthy group. Similarly, it also ranks lower in samples from Canada compared to those from Ireland (**Supplementary Fig. 2**). The pre-trained MGM model outperformed the un-pre-trained model in both cross-regional and local region diagnostics. After partial fine-tuning with data from the target region, the cross-regional diagnostic performance reached satisfactory levels (Ireland predicting Canada, ROC-AUC: 0.844; Canada predicting Ireland, ROC-AUC: 0.829, **Fig. 4a, 4b and 4c**). For the irritable bowel syndrome (IBS) cohort [36] consisting of samples from Australia, the United Kingdom, and the United States, the model’s initial performance was poor in Australia due to small sample sizes (n=21, ROC-AUC: 0.500). However, after training on a larger dataset from the United Kingdom (n=171) or United States (n=498) and fine-tuning with data from the target countries, the model’s diagnostic performance in Australia and the United Kingdom improved significantly (Australia from United Kingdom ROC-AUC: 0.833; United Kingdom from United State ROC-AUC: 0.800, **Fig. 4d, 4e and 4f).** Our results aligned with previous studies based on fully connected neural network [37], suggested building a foundation diagnostic model on a large, diverse dataset and then fine-tuning it with smaller regional datasets proves to be an effective strategy for mitigating regional effects.

**Figure 4.**
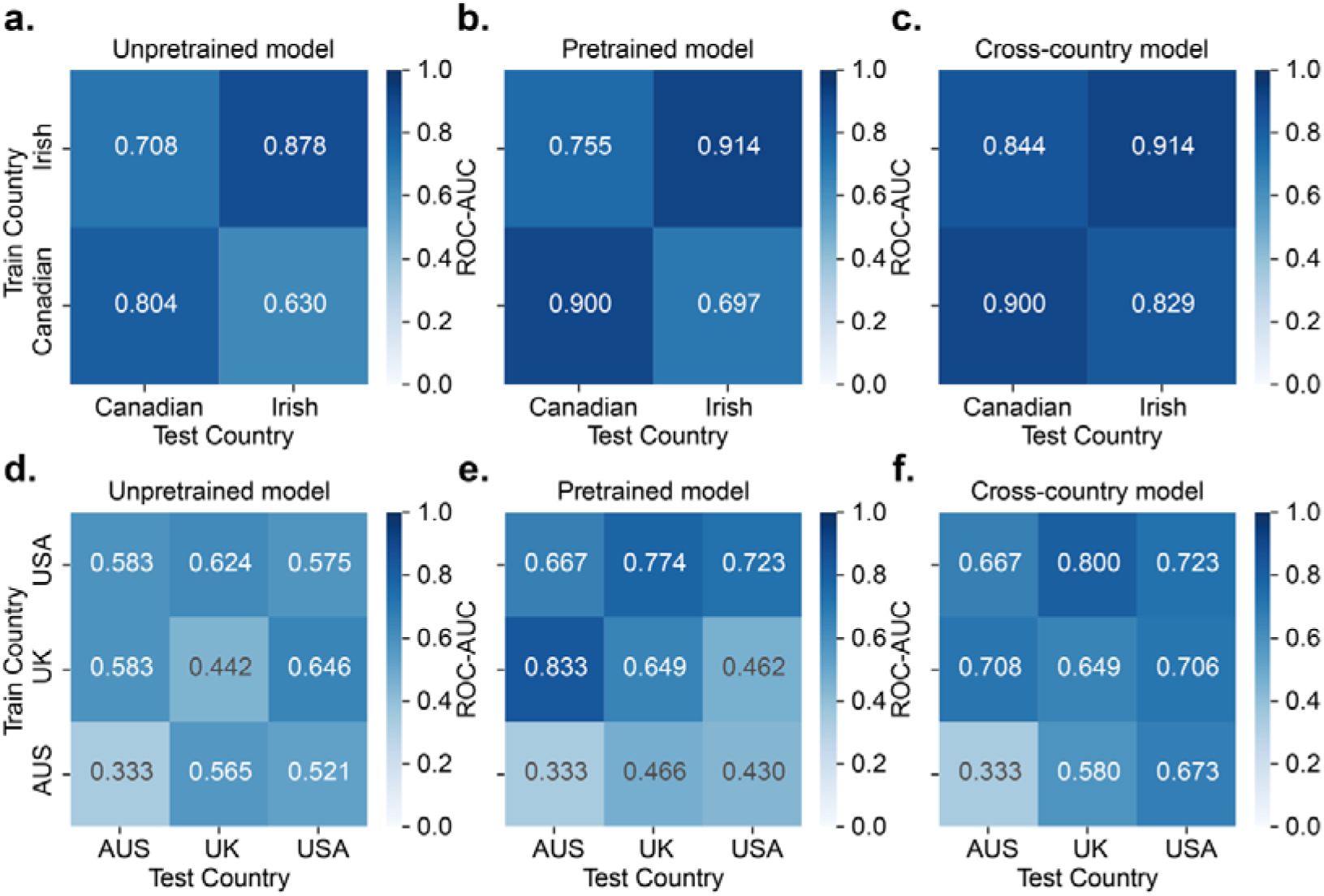
Result of disease diagnosis across intercontinental regions. **a.** Heatmap of ROC-AUC of un-pre-trained model on IBD cohort. **b.** Heatmap of ROC-AUC of pre-trained model on IBD cohort. **c.** Heatmap of ROC-AUC of cross-country model on IBD cohort. **d.** Heatmap of ROC-AUC of un-pre-trained model on IBS cohort. **e.** Heatmap of ROC-AUC of pre-trained model on IBS cohort. **f.** Heatmap of ROC-AUC of cross-country model on IBS cohort. Row represents countries for training the model. Columns represents countries for evaluating the model.

### Infant development monitoring and keystone genus discovery

Next, we fine-tuned our model on a longitudinal infant dataset from Roswall et al. [38], to distinguish the developmental stages of infants from different delivery modes (**Fig. 5a**). During infancy, the gut microbiome undergoes dynamic changes and typically stabilizes in childhood. Meanwhile, the mode of delivery significantly impacts the microbiome in neonates, which can be linked to health outcomes later in life [39]. However, relatively few methods exist to uncover the dynamics progression of gut microbiota from infancy toward a stable, adult-like microbiota.

**Figure 5.**
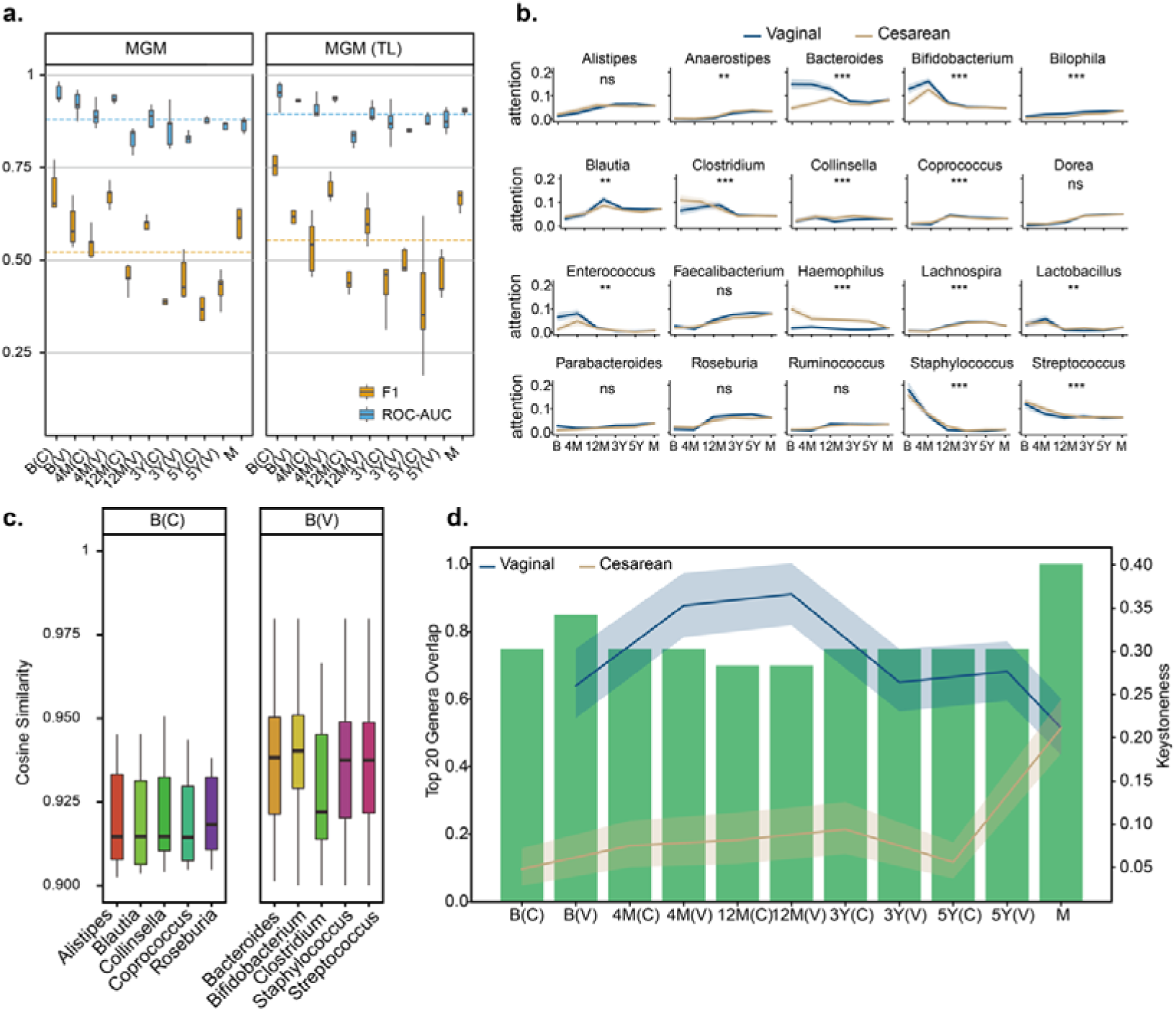
MGM Enhanced age prediction and keystone genus discovery. **a.** Boxplot evaluation of MGM model using 5-fold cross-validation, with blue representing ROC-AUC, orange representing F1, and dashed lines indicating the average of all experiments. **b.** Line plot showing the attention weights of the top 20 genera across layers and heads. Asterisks representing the significance of Kolmogorov–Smirnov (KS) test: ***: *P* < 0.005, **: *P* < 0.01, *: *P* < 0.05, ns: not significant. **c.** Top 5 genera with deleterious effects at birth from cesarean deliveries and birth from vaginal deliveries. **d.** Keystoneness of genera appearing in at least 30% of samples in each group (pink line plot). Overlap of the top 20 highest attention weight genera and the top 20 highest keystoneness genera (green bar plot). B: birth, 4M: 4 months, 12M: 12 months, 3Y: 3 years, 5Y: 5 years, C: cesarean, V: vaginal, M: mother, TL: transfer learning.

Our model could accurately predict the developmental stage and delivery mode of these infants, outperforming both MGM trained from scratch and traditional methods. In comparison, EXPERT, a supervised pre-training method based on neural network for microbial community classification [22], performed poorly in this task, indicating the superiority of self-supervised pre-training methods and attention mechanisms (EXPERT, average ROC-AUC = 0.54; EXPERT with transfer learning, average ROC-AUC = 0.54; MGM, average ROC-AUC = 0.90; MGM with transfer learning, average ROC-AUC = 0.91, **Fig. 5a**, **Supplementary Fig. 3a**). Besides, we visualized sample embeddings to capture the dynamic changes in microbial communities at different stages (**Supplementary Fig. 4**). The high similarity between samples of cesarean delivery and vaginal delivery samples at the same development stage suggest that infant gut microbes are stage-specific. As the infant ages, the sample embeddings become increasingly similar to their mother’s, indicating that infant gut microbial communities evolve toward those of adults.

We then examined the attention weights from 64 attention across developmental stages. Several genera, including *Dorea*, *Faecalibacterium* and *Ruminococcus* (KS test, *P* = 0.13, 0.19 and 0.22), demonstrated similar attention trends across delivery modes, consistent with their known associations with infant age [40]. However, most genera demonstrated distinct attention patterns. Notably, *Bacteroides* (KS test, *P =* 5.87E-5), a keystone taxon of the human gut microbiota [41] and *Bifidobacterium* (KS test, *P =* 1.53E-12), a common probiotic [42], received higher attention weights in vaginal deliveries during early stages. In contrast, *Haemophilus* (KS test, *P =* 4.88E-90), a well-known human pathogen, had consistently higher attention weights in cesarean deliveries throughout the entire developmental process (**Fig. 5b**).

We further employed a leave-one-genus-out deletion approach to identify genera whose removal would have a deleterious effect in this context. In this analysis, one genus was removed from the rank value encoding at a time, and the impact on the embeddings of the remaining genera was quantified by similarity. The deleterious pattern of two delivery modes is consistent with the results of the attention weights analysis that probiotic has higher deleterious effects on infants delivered vaginally. Keystone taxa such as *Bacteroides*, *Roseburia*, and *Faecalibacterium* exhibited the higher deleterious effects (**Fig. 5c and Supplementary Fig. 3b**). Based on these findings, we hypothesize that genera with high attention weights and deleterious effects possess strong keystone attributes.

We quantified the keystone attributes of genera using DKI framework [43], a deep-learning model designed to assess community-specific keystoneness by conducting thought experiment on species removal. We found that genera with high attention weight had a large overlap with genera with high keystoneness, and these genera were present in a large proportion of samples. Interestingly, infants delivered vaginally exhibited higher overall keystoneness compared to those delivered by cesarean section, with the mother’s keystoneness was intermediate between the two delivery modes. (**Fig. 5d and Supplementary Fig. 3c**). In summary, several bacterial taxa identified as keystone in gut microbiome exhibited high attention weights in our model. This demonstrated that our attention-based model can effectively capture keystones taxa and the dynamic developmental trajectory of microbial community, paving the way for comprehensive analysis of diverse microbiome from both spatial and temporal perspectives.

### Potential cancer treatment target identification

We next sought to explore the potential applications of MGM model in distinguishing different types of cancers and identifying tumor-specific biomarkers. Recent studies have increasingly highlighted the presence of microbial signals within tumor tissues, suggesting that the tumor microbiome could be a valuable target for cancer research and treatment [44–46]. However, the research on biomarkers across different tumor tissues may not yet be sufficiently comprehensive [47]. Additionally, the ability to accurately diagnose and identify multiple types of tumor tissues remains limited in precision. This underscores the need for more robust models that can effectively differentiate cancers and identify specific microbial biomarkers, which could be crucial for developing targeted therapies.

To this end, we fine-tuned our model using five types of gastrointestinal tumors obtained from The Cancer Microbiome Atlas (TCMA) database [48]. Firstly, our model with a macro-ROC value reaching 0.97 (**Fig. 6a**). Specifically, the ROC values for COAD, ESCA, HNSC, READ, and STAD were 0.99, 0.97, 0.98, 0.98, and 0.97, respectively, indicating a robust performance across different cancer types.

**Figure 6.**
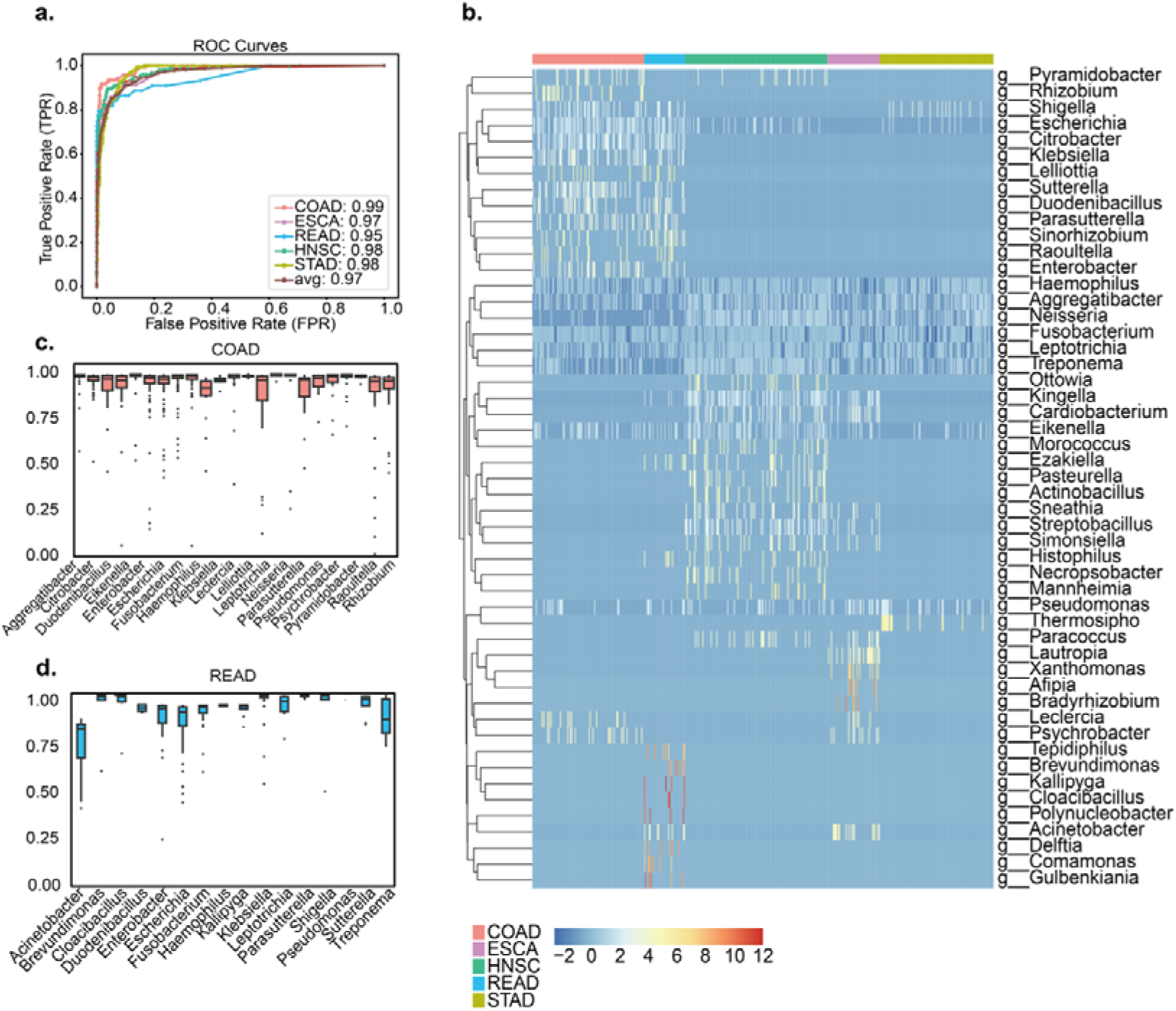
MGM Enhanced Cancer Diagnosis and Biomarker Identification. **a.** ROC performance of MGM in distinguishing between five types of cancer. The x-axis represents the False Positive Rate (FPR), while the y-axis indicates the True Positive Rate (TPR). The macro-average ROC value reaches 0.97, with individual ROC values as follows: COAD (0.99), ESCA (0.97), HNSC (0.98), READ (0.98), and STAD (0.97). **b.** Distribution of the top 50 species identified by attention across the five types of cancer. The x-axis represents different cancer types (COAD, ESCA, HNSC, READ, STAD), and the y-axis lists the microbial genera. The color intensity in each cell corresponds to the abundance level, with the scale ranging from -2 to 12, where positive values indicate a higher relative abundance. **c-d.** Cosine similarity of embedding vectors before and after the removal of a specific genus in COAD and READ. The x-axis shows the microbial genera removed, and the y-axis represents the cosine similarity score, ranging from 0 to 1.

Secondly, to further explore the potential therapeutic targets, we employed the ‘leave-one-genus-out’ approach to assess the impact of each genus’s absence on the sample embeddings. We extracted the top 50 biomarkers ranked by MGM-calculated attention scores, and the heatmap (**Fig. 6b**) illustrated their distribution across the five cancer tissues. Our findings suggested that some genera exhibited significant abundance differences in particular types of cancer. For instance, *Escherichia* and *Enterobacter* had a significant detrimental impact on COAD and READ samples. These genera have been previously associated with gastrointestinal tumors [48–50]. In COAD samples, the abundance of *Escherichia* was observed to be 7.25 times higher than in non-COAD samples. Conversely, genera like *Acinetobacter* may act as key biomarkers distinguishing COAD and READ tissues. *Streptobacillus*, another significant genus, displayed an elevated presence in HNSC, with a relative abundance increase of 5.26 times compared to other types, indicating its importance in distinguishing HNSC. Boxplots (**Fig. 6c-d**) showed the cosine similarity of the embedding vectors before and after the removal of specific genus. Notably, when *Acinetobacter* was removed, the cosine similarity in the READ samples significantly decreased, underscoring the critical role this genus plays in shaping the microbial composition of these samples. These findings not only validated the effectiveness of MGM model in identifying cancer-associated microbial biomarkers, but also offered new possibilities for targeted cancer diagnosis and treatment.

Collectively, our model demonstrated high diagnostic accuracy and robust performance across multiple cancer types, underscoring the crucial role of the tumor microbiome in cancer development. These advancements revealed MGM’s potential for more precise and targeted cancer therapies, highlighting the potential of integrating tumor microbiome analysis into clinical practice.

### In silico perturbation analysis and validation on Crohn’s disease

Microbial perturbation studies are crucial for understanding the impact of treatments, such as antibiotics, on microbiome composition in disease contexts. To verify MGM’s capability to capture microbial perturbations, we fine-tuned the model on a dataset containing intestinal mucosa microbiome samples from Crohn’s disease (CD) patients before and after antibiotic treatment [51].

In our in silico perturbation analysis, we identified microbes whose enrichment or reduction in CD patients’ intestines could shift sample embeddings towards those of healthy controls. To simulate microbial enrichment, we repositioned a genus token to the next position after the ‘bos’ token, while microbial reduction was simulated by repositioning the genus token to the position before the ‘eos’ token. Our results aligned with the original study, particularly in CD patients who had not received antibiotic treatment (**Fig. 7a-d**). In both Terminal ileum and Rectum samples, the in silico reduction of *Enterobacteriaceae*, *Pasteurellaceae*, *Veillonellaceae*, *Fusobacteriaceae*, and *Neisseriaceae* resulted in a higher similarity to healthy controls compared to in silico enrichment, mirroring findings from the original study. However, *Gemellaceae*, another family reported as increased in the original study, showed no difference between in silico enrichment and reduction.

**Figure 7.**
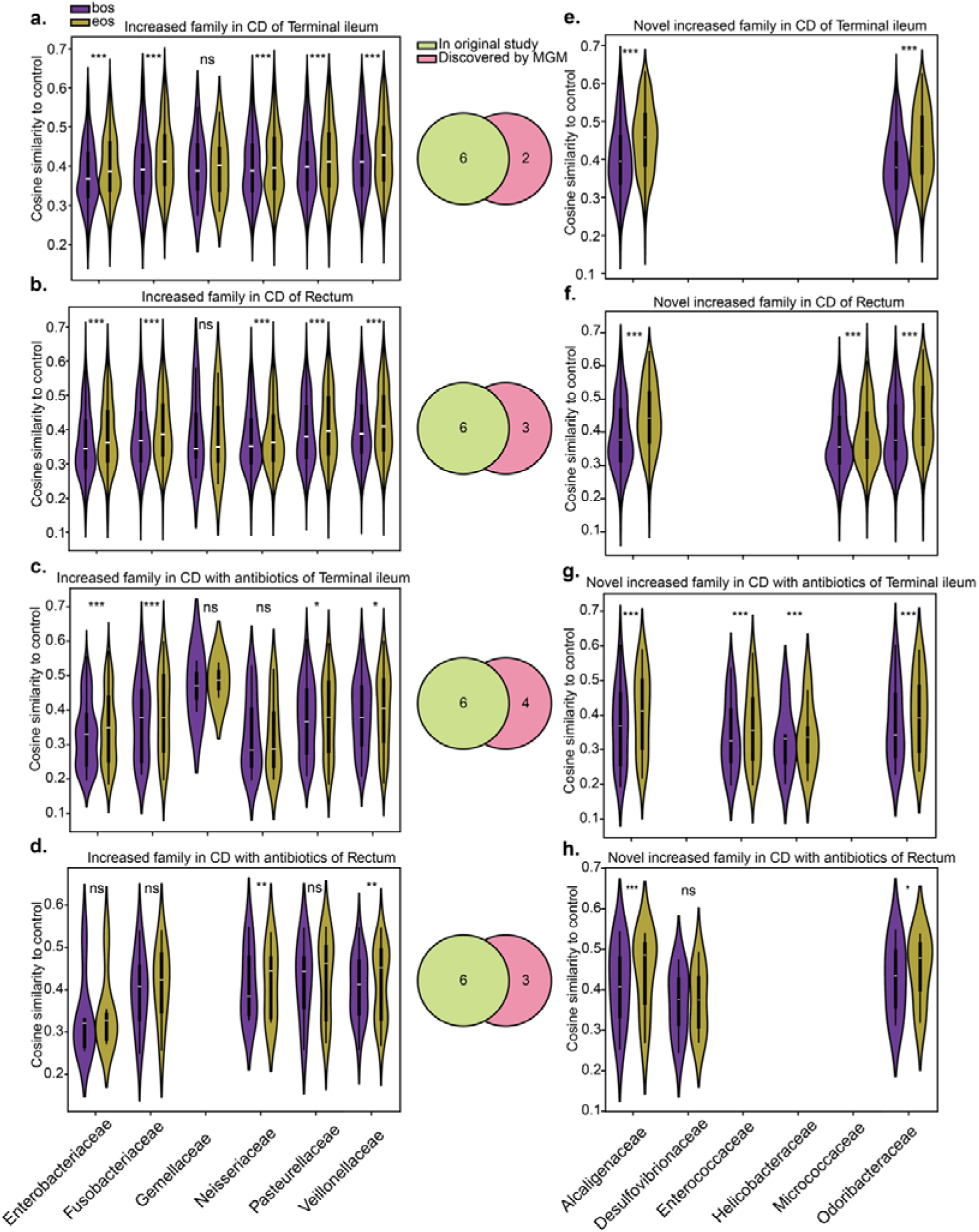
In silico enrichment and reduction analysis on CD dataset. **a.** In silico analysis of six families reported as increased in the Terminal ileum of CD patients. **b.** In silico analysis of six families reported as increased in the Rectum of CD patients. **c.** In silico analysis of six families reported as increased in the Terminal ileum of CD patients treated with antibiotics. **d.** In silico analysis of six families reported as increased in the Rectum of CD patients treated with antibiotics. **e.** In silico analysis of novel families identified as increased in the Terminal ileum of CD patients. **f.** In silico analysis of novel families identified as increased in the Rectum of CD patients. **g.** In silico analysis of novel families identified as increased in the Terminal ileum of CD patients treated with antibiotics. **h.** In silico analysis of novel families identified as increased in the Rectum of CD patients treated with antibiotics. ‘bos’ indicates enrichment simulation by shifting a genus token to the next position after the ‘bos’ token, while ‘eos’ indicates reduction simulation by shifting a genus token to the position before the ‘eos’ token. The Y-axis represents the cosine similarity of the sample embedding to that of healthy controls. Venn diagrams in the center show the overlap between families reported as increased in the original study and the top six families showing the largest differences in our in silico analysis, as well as novel families identified through this analysis.

We further evaluated the top six families showing the largest differences in our in silico analysis against the six families reported as increased in the original study. We identified several novel families that were more prevalent in CD patients, including *Alcaligenaceae* and *[Odoribacteraceae]*, whose in silico reduction significantly increased similarity to healthy controls (**Fig. 7e-h**). Previous studies have shown that *Alcaligenaceae* is enriched in mesenteric adipose tissue in CD patients [52], and *[Odoribacteraceae]* is more abundant in CD patients with the Type II Paneth cell phenotype [53]. Moreover, the in silico reduction of *Enterococcaceae* and *Helicobacteraceae* in Terminal ileum samples from CD patients treated with antibiotics exhibited a significantly greater impact, suggesting that these families may show resistance to antibiotic treatment (**Fig. 7g**).

These findings highlighted MGM’s effectiveness in detecting both known and novel microbial perturbations. By capturing microbial dynamics that align with experimental data and identifying new perturbations, MGM proved its utility for microbiome research, particularly for exploring therapeutic impacts in disease contexts.

## Discussion

In this study, we proposed MGM, the first foundation model designed for microbial community analysis, leveraging pre-trained transformers on a diverse corpus of microbiome data. By employing large-scale self-supervised pre-training, MGM develops a foundational understanding of microbial interactions within communities, free from task-specific biases. This general representation captures broad patterns and relationships across varied microbiome datasets, establishing MGM as a versatile tool in microbiome research.

Benchmark evaluations underscore MGM’s superior performance in microbial community classification tasks. In cross-validation on the Microcorpus-260K dataset, the fine-tuned MGM achieved an average ROC-AUC of 0.99, significantly outperforming traditional methods, including source tracking techniques and machine learning models. Its application to 43,528 additional samples from MGnify revealed exceptional performance in deeper, more complex analyses. Furthermore, MGM embeddings enabled seamless integration of microbiome data across different sources and batches, highlighting its utility in distinguishing microbial samples for tasks such as microbial source tracking.

To tailor these general insights to specific microbiome-related tasks, MGM employs a contextualization approach. By fine-tuning the foundation model on task-specific datasets, MGM adapts its learned representations to align with the unique characteristics and nuances of the target task. This process enhances the model’s performance on various microbiome-related tasks by leveraging the robust, broad knowledge acquired during pre-training. Through systematically analyzing the performance of MGM, we gain valuable insights into how pre-trained language model can be applied to microbiome data analysis, showing MGM’s potential for enhancing performance on various microbiome-related tasks.

Beyond classification, MGM captures the spatial and temporal dynamics of microbial communities. In cross-regional intestinal disease datasets, MGM overcame regional limitations, achieving accurate diagnoses across intercontinental regions. When applied to a longitudinal infant dataset, the model effectively traced the development and maturation of the infant gut microbiome. Attention-weight analyses across developmental stages and delivery modes identified key genera, such as *Bacteroides* and *Bifidobacterium*, which were more prominent in vaginal deliveries, while *Haemophilus* showed higher weights in cesarean deliveries.

In silico perturbation experiments further highlighted MGM’s clinical potential. Fine-tuning on the TCMA database revealed genera with significant deleterious effects across various tumor types, suggesting its utility in identifying microbial targets for cancer therapy. When applied to a Crohn’s disease dataset involving antibiotic treatment, MGM detected microbial perturbations in the intestinal mucosa consistent with the original study, along with novel findings, such as shifts involving *Alcaligenaceae* and *Odoribacteraceae*, later corroborated by independent research. These findings illustrate MGM’s sensitivity to microbial community changes and its potential for therapeutic applications.

While MGM model shows promise, several limitations need to be addressed. The primary limitation lies in its rank value encoding method. While this method effectively mitigates the impact of extreme values and converts tabular data into sequential data, it fails to preserve the original relative abundance information. This shortcoming complicates the reconstruction of samples into their original abundance tables, limiting the model’s generative capabilities. Future work should focus on refining the encoding process to better retain relative abundance information. Another area for improvement involves the expansion of the model’s training dataset. While MGM has demonstrated strong performance, its generalizability could be further enhanced by incorporating a broader range of microbiome samples from different biomes and populations. Given the model’s ease of fine-tuning, updating it with additional datasets is a practical way to boost its adaptability across various microbiome-related tasks. Additionally, while the model shows great promise in identifying treatment targets and keystone genera, incorporating wet-lab experimental validation would strengthen the robustness and comprehensiveness of our findings.

In conclusion, MGM represents a profound advancement in microbiome research, offering a robust and adaptable tool for analyzing microbial communities. As a foundation model, MGM exceled in large-scale microbial classification, leveraging vast pre-trained datasets to capture fundamental patterns that span diverse microbial ecosystems. In its contextualized form, MGM could be fine-tuned for specific downstream tasks, such as identifying clinically relevant microbial perturbations and uncovering nuanced microbial interactions. This dual capacity to model both general and task-specific patterns underscored its broad applicability in microbiome science, including therapeutic interventions and diagnostic innovations. As a powerful foundation model, MGM paves the way for future innovations in microbiome analysis, contributing to a deeper understanding of microbial ecosystems and their roles in human health.

## Methods

### Data Preprocessing

We assembled a comprehensive dataset, Microcorpus-260K, which includes all samples from MGnify up to June 2023. Initial processing involved retaining genus-level relative abundances and filtering out genera with relative abundances less than 0.01%. Samples were further filtered to retain only those with at least 10 genera with non-negligible abundance, resulting in a final dataset of 263,302 samples and a vocabulary of 9,665 genera.

For each sample, we standardized the relative abundances of each genus using their mean and standard deviation across all samples. These standardized values were then rank encoded to prepare the data for model input. The means and standard deviations calculated during this step were saved for future standardization of downstream data.

### Model Architecture

We constructed MGM model using eight layers of transformer blocks, with each block consisting of a self-attention layer and a feed forward neural network layer. Given the fixed input length requirement of transformer models, we set the input length to 512 tokens, covering 99.99% of the samples. This length ensured that most samples could be processed without truncation, preserving the integrity of the data. Additional key hyperparameters were as follows: activation function, Gaussian Error Linear Unit (GELU); attention heads per layer, eight; feed forward size, 1024. The modeling framework was implemented in PyTorch, leveraging the Huggingface Transformers library for model configuration and training [54].

The self-attention mechanism employed in each transformer layer follows the scaled dot-product attention formula:

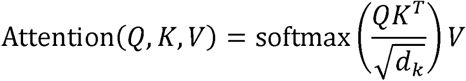

Where *Q* (queries), *K* (keys), and *V* (values) are the linear projections of the input, and *d_k_* is the dimensionality of the keys. This formulation allows the model to capture contextual relationships between different tokens in the sequence, enabling more effective representation learning for downstream tasks.

### Pre-training Procedure

Pre-training was conducted using the causal language modeling approach with self-attention mechanism to capture co-occurrence pattens among tokens. For each sample, we appended a ‘bos’ token at the beginning and an ‘eos’ token at the end to denote the start and end of a sequence, respectively. Different from transformer encoder models like BERT, which randomly masked some tokens and predicted them in the output, our autoregressive model was trained to predict the next possible token in the sequence based on known input tokens, facilitating the learning of contextual relationships among genera within a sample. Specifically, the order of predicted tokens also implied genera relative abundance distribution in the sample.

Training was executed using Huggingface’s Trainer API. Key hyperparameters included: learning rate, 1e-3; batch size, 50; warmup steps, 1000; weight decay, 0.001; validation split, 10% of the data. Validation loss is calculated per 500 training steps, Early stopping based on validation loss with 3 patience.

### Model interpretability

We conducted an interpretability analysis by leveraging the attention weights extracted from the multi-head, multi-layer transformer. These attention weights were modified by replacing *V* with *V*^0^, where *V*^0^ represents one-hot indicators for each position index. To consolidate the attention information across the model, we integrated the attention matrices by calculating an element-wise average across all layers and attention heads. To identify the genera with the highest attention weights in a microbial community, we summed the attention weights across each column, obtaining the total attention weight from a single genus to all other genera in the community.

### Sample Representation

Each sample is analogous to a ‘sentence’ composed of genera, and its representation is obtained by aggregating the learned genus-level representations. In this study, we opted element-wise mean pooling to get the sample representation from our pre-trained model. For fine-tuned model, as the last token (‘eos’ in this study) was used to do the sequence classification, we used its embeddings as the sample representation.

### Downstream Fine-tuning

For downstream tasks, the pre-trained MGM model is fine-tuned by replacing the language modeling head with a task-specific head. All downstream tasks in this study focused on microbial community classification. Fine-tuning employed a sequence classification head, which utilized the final token (‘eos’ in this study) for classification. Fine-tuning was executed using Huggingface’s Trainer API. Key hyperparameters included, learning rate, 1e-3, batch size, 50, warmup steps, 1000, weight decay, 0.001, validation split, 10% of the data. For microbial classification task on MicroCorpus-260K, For the microbial classification task on the MicroCorpus-260K dataset, validation loss was calculated every 500 training steps, while for other downstream tasks, it was calculated per training epoch. Early stopping based on validation loss with 3 patience.

The evaluation of both the microbial classification task and the infant age prediction task was performed using a 5-fold cross-validation strategy. Training was conducted on 80% of the samples, with performance tested on the 20% held-out samples, and this process was repeated across five folds. For the cross-regional disease diagnosis task, 50% of the samples from each region were split as the test set using stratified sampling, while the remaining 50% were used either to train the diagnostic model or fine-tune a model trained on another region. Notably, the fine-tuning applications were trained on classification objectives distinct from the causal language modeling objective, making the inclusion of task-specific data in the pre-training corpus irrelevant to classification predictions.

### Comparison methods

**FEAST:** FEAST is an expectation-maximization-based method that estimates the proportion of the sink community contributed by various source environments [34]. For benchmarking, we employed the R package implementation of FEAST.

**EXPERT:** EXPERT is an ontology-aware neural network method that leverages transfer learning for microbial community classification [22]. We benchmarked EXPERT using its Python package.

**DKI:** DKI is a deep learning-based approach designed to identify keystone species in microbial communities [43]. We utilized the scripts provided in DKI’s GitHub repository to validate the keystone microbes identified by MGM.

**Other Machine Learning Methods:** Additional machine learning models used for benchmarking, including K-Nearest Neighbor, Logistic Regression, and Random Forest, were implemented using the scikit-learn library [55].

### Code Availability

The code for MGM model is available at https://github.com/HUST-NingKang-Lab/MGM.

## Acknowledgments

This work was partially supported by the National Key R&D Program of China (Grant No. 2023YFA1800900 and 2018YFC0910502), the National Natural Science Foundation of China (Grant Nos. 32071465, 31871334, 81827901). Numerical computations were performed on the Hefei Advanced Computing Center.

## Author contributions

HZ and KN conceived of and proposed the idea, designed and developed the framework. HZ and YZ performed the experiments and analyzed the data. HZ, YZ and ZK visualized the data. HZ, YZ, ZK, LS and KN contributed to editing and proof-reading the manuscript. All authors read and approved the final manuscript.

## Competing interest

The authors declare that they have no competing interests.

## Ethics approval and consent to participate

Not applicable.

## Notes

https://github.com/HUST-NingKang-Lab/MGM

### Competing Interest Statement

The authors have declared no competing interest.

