## Supplementary material for "MGM as a large-scale pretrained foundation model for microbiome analyses in diverse contexts"

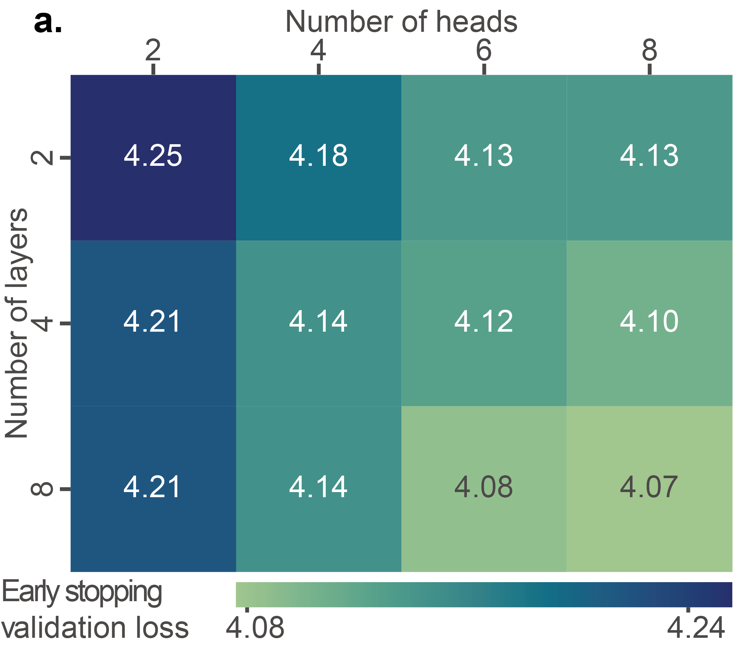

**Supplementary Figure 1. Early stopping validation loss across different grid search configurations.** Rows indicate the number of transformer layers, and columns represent the number of attention heads per layer. During pre-training, the batch size was set to 64, and validation loss was monitored every 500 training steps. Pre-training was halted if the validation loss failed to decrease for five consecutive evaluations.

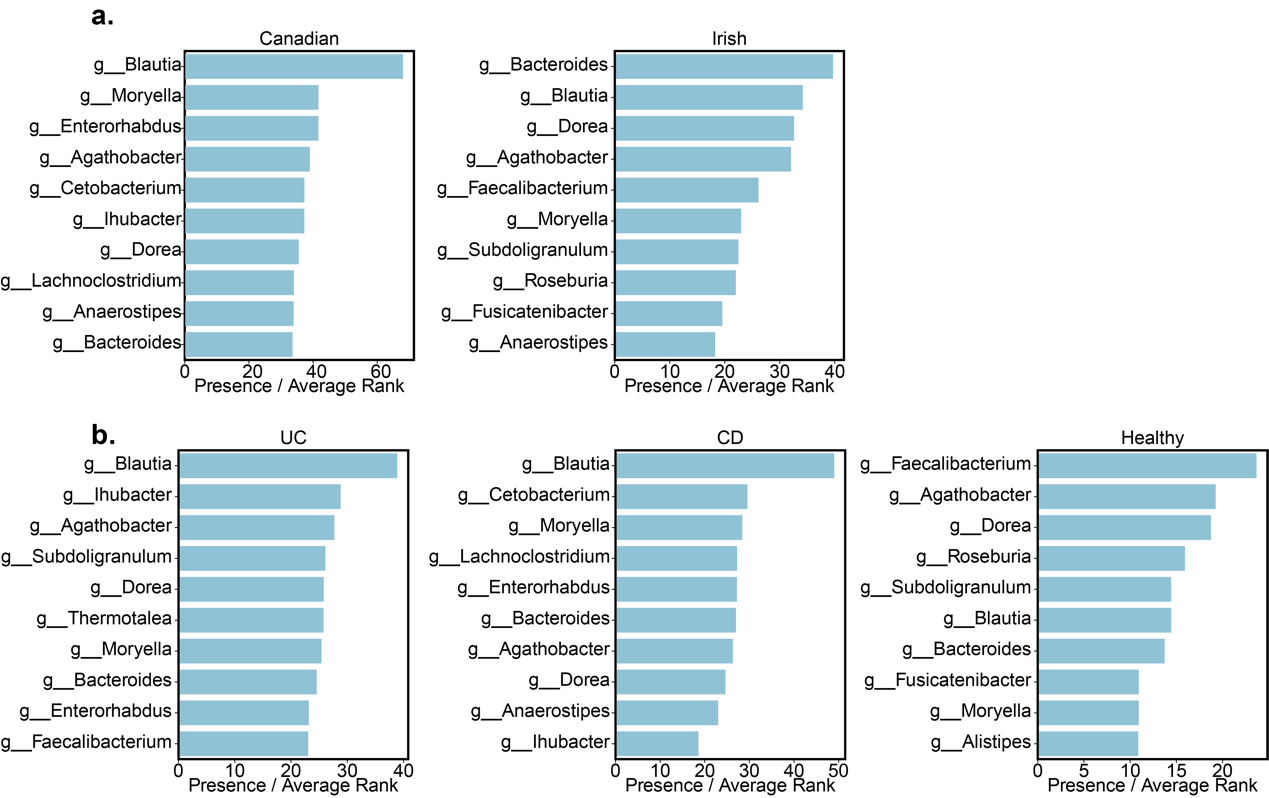

**Supplementary Figure 2. Candidate genus of regions and diseases of IBD cohort. a.** Top 10 Candidate genera of different regions. **b.** Top 10 Candidate genera of different diseases. Candidate genera are measured by their number of occurrences divided by the average rank in each group.

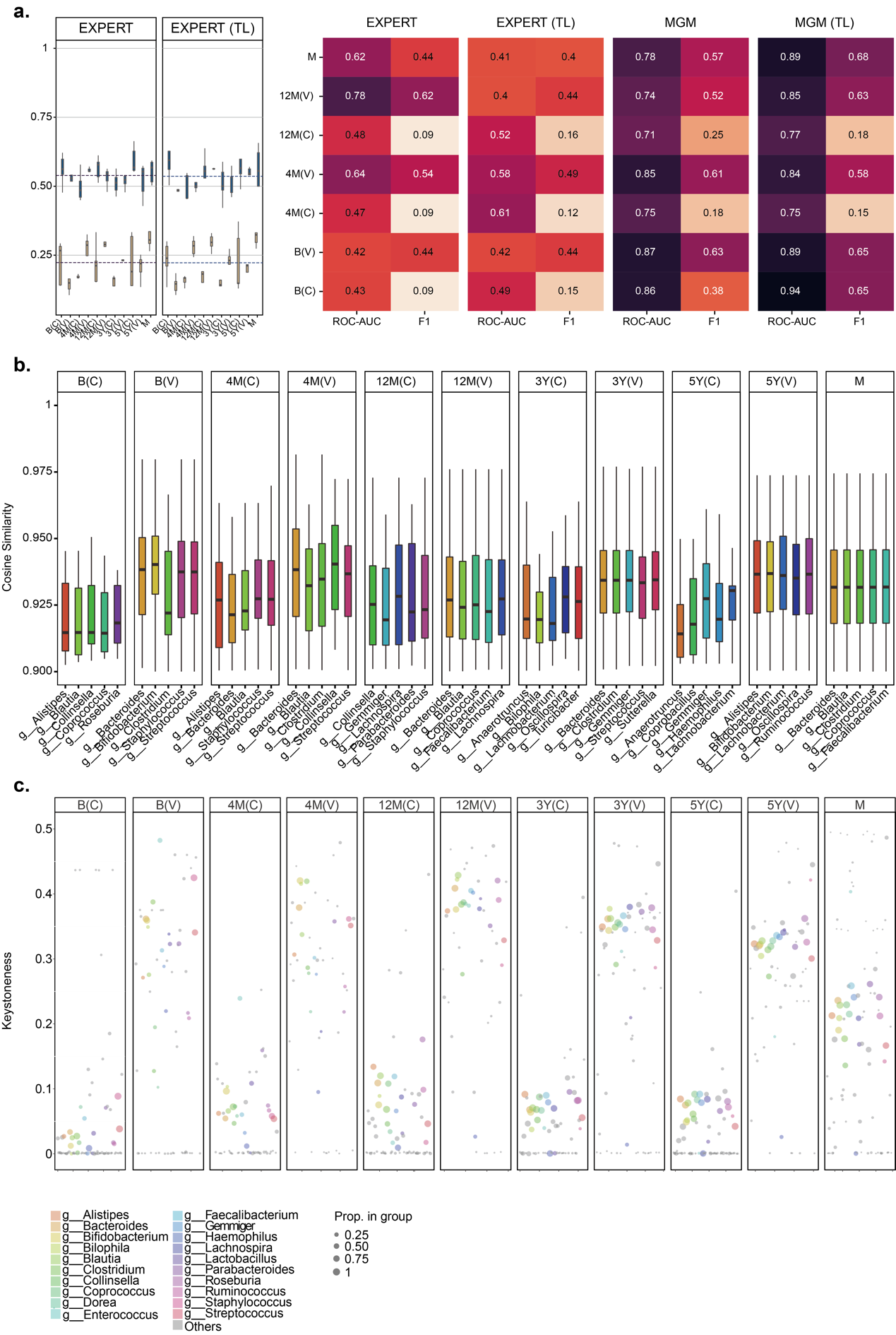

**Supplementary Figure 3. Evaluation on infant datasets and excavation of keystone genera. a.** EXPERT model performance and comparation with MGM on infant dataset. **b.** Top 5 genera with highest deleterious effects in each development stage. **c.** Keystoness of genera with highest attention weights. Top 20 genera with highest attention weights are colored. The size of point represents the proportion that occurs in each development stage.

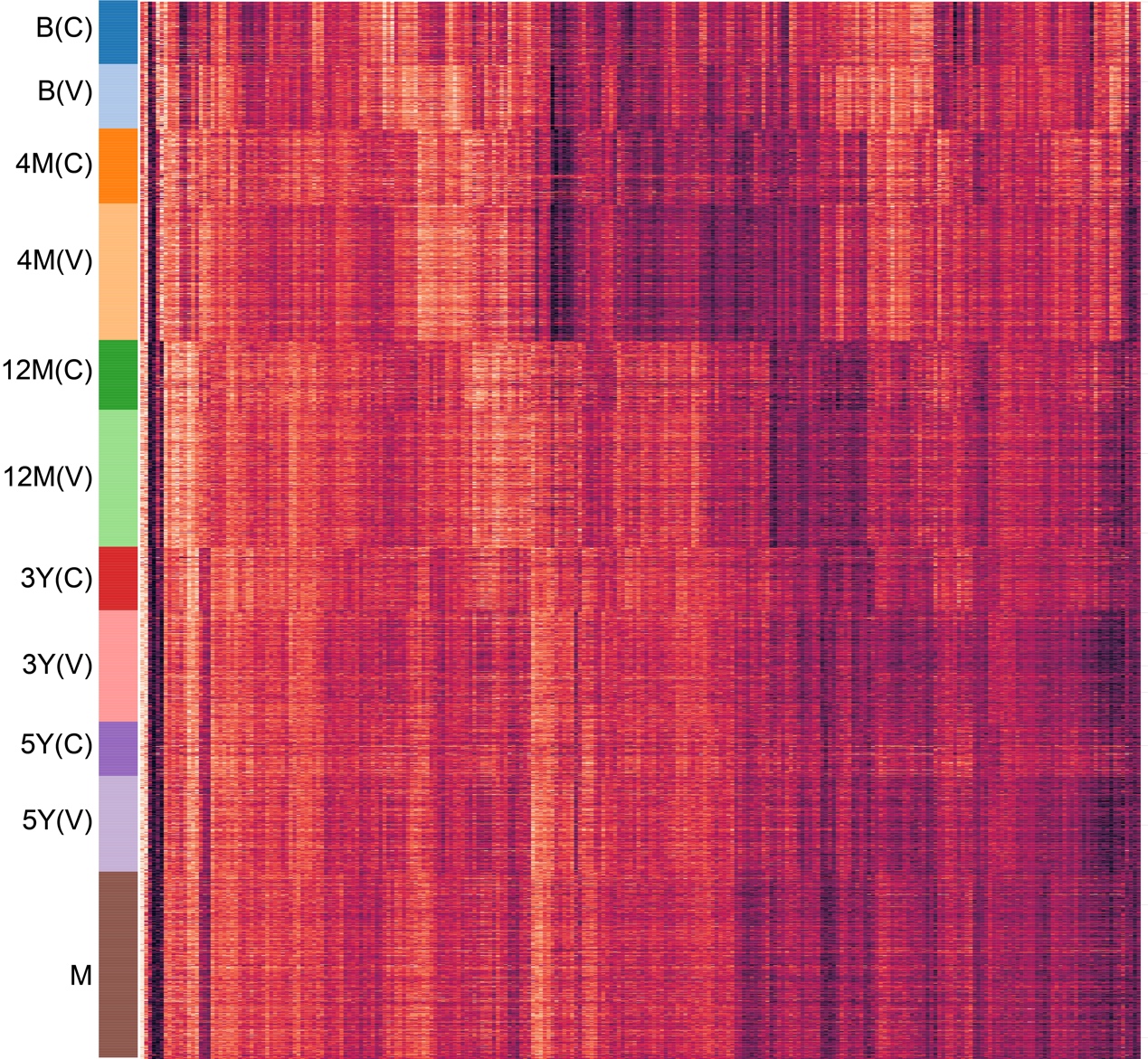

**Supplementary Figure 4. Heatmap of 256-dimentional sample embeddings from MGM in infant dataset.** Rows represent sample and ordered by development stage and delivery mode. Columns represent the embedding value of each dimension. Clustered by columns.
